# SPUMONI 2: Improved pangenome classification using a compressed index of minimizer digests

**DOI:** 10.1101/2022.09.08.506805

**Authors:** Omar Ahmed, Massimiliano Rossi, Travis Gagie, Christina Boucher, Ben Langmead

## Abstract

Genomics analyses often use a large sequence collection as a reference, like a pangenome or taxonomic database. We previously described SPUMONI, which performs binary classification of nanopore reads using pangenomic matching statistics. Here we describe SPUMONI 2, an improved version that is faster, more memory efficient, works effectively for both short and long reads, and can solve multi-class classification problems with the aid of a novel sampled document array structure. By incorporating minimizers, SPUMONI 2 reduces index size by a factor of 2 compared to SPUMONI, yielding an index more than 65 times smaller than minimap2’s for a mock community pangenome. SPUMONI 2 also achieves a speed improvement of 3-fold compared to SPUMONI and 15-fold compared to minimap2. We show SPUMONI 2 achieves an advantageous mix of accuracy and efficiency for short and long reads, including in an adaptive sampling scenario. We further demonstrate that SPUMONI 2 can detect contaminated contigs in genome assemblies, and can perform multi-class metagenomic read classification.

## 1 Introduction

Read classification is a component of many sequencing data analyses, such as taxonomic classification (13, 20, 34), host sequence depletion (16, 18), and adaptive sampling of nanopore reads (14, 26). Databases holding reference sequences are growing rapidly (19, 30), enabling pangenomic methods that use a collection of related genomes as the reference, rather than a single genome. We previously described SPUMONI (1), a method for rapid binary classification of nanopore reads against a pangenome reference. SPUMONI builds on the *r*-index (9, 15), a compressed index that grows with the amount of distinct sequence in the reference pangenome. It uses the MONI algorithm (29) to compute *matching statistics* (defined below) as input to its classification decision. Though SPUMONI was faster and used less memory than a minimap2-based approach, we since sought ways to extend its functionality to (a) analyze short as well as long reads, (b) perform multi-class as well as binary classification, and (c) scale efficiently to larger pangenomes.

We present SPUMONI 2, which uses a minimizer scheme to digest and reduce both the pangenome reference and the input reads. By changing the alphabet to consist of all possible minimizers, rather than all possible bases, SPUMONI 2 queries are faster than SPUMONI’s. Moreover, the approach of SPUMONI 2 works in any situation where a read that truly originates from the reference pangenome yields longer exact substring matches than a read that does not. SPUMONI 2 works with long or short reads and does not require the user to have foreknowledge of what *k*-mer length is capable of distinguishing true from false hits.

Both SPUMONI and SPUMONI 2 build upon the MONI algorithm (29) for computing matching statistics. The *matching statistics* of a pattern string with respect to a collection of reference strings is an array storing the length of the longest prefix of the pattern’s *i*^th^ suffix that occurs in the reference. For faster queries, SPUMONI 1 & 2 compute an approximation of matching statistics called *pseudomatching lengths* (PMLs). See Methods (Section 4) for formal definitions.

Minimizers (28) are a form of locality-sensitive hashing, applied to windows of a string. A minimizer scheme defines a small window size (*k*) and a large window size (*w*); within each length-*w* window, the constituent *k*-mer with the minimal hash value is chosen as the minimizer. They have been used to create small genomic and pangenomic indexes (18, 34). In some settings, minimizers are used to *digest* a long sequence, replacing it with a concatenation of its minimizers (with equal-minimizer runs collapsed). This can, for instance, reduce the size of assembly graph representations (7, 8).

SPUMONI 2 first creates a minimizer digest with configurable small (*k*) and large (*w*) window sizes, then builds the *r*-index over the digest. When indexing the digest, SPUMONI 2 considers the alphabet to consist of all possible choices of minimizer (i.e. all possible *k*-mers) (7, 8). That is, it uses a minimizer alphabet rather than a DNA alphabet. This poses no problem to the *r*-index, which can adapt to any discrete alphabet. This combined strategy of indexing the minimizer digest and using a minimizer alphabet (see Figure 1), has the effect of reducing index size, memory footprint, and query time.

**Figure 1:**
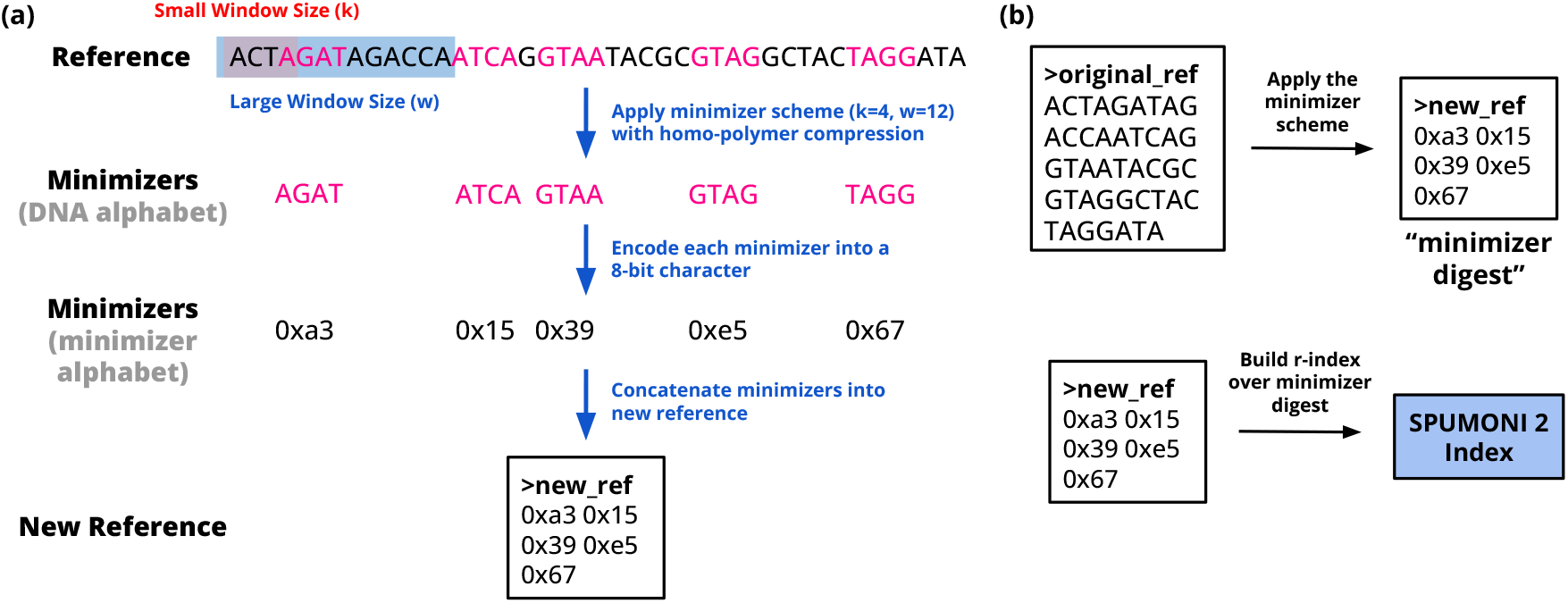
(a) Shows the procedure used by SPUMONI 2 to digest a reference into a smaller reference of concatenated minimizers. The *k*-mer that minimizes a hash function in each *w*-bp window as well as being distinct from the previous minimizer is chosen and represented as an 8-bit character in the new reference. In practice, SPUMONI 2 would also include the reverse complement of each sequence prior to applying the minimizer scheme. (b) After generating this minimizer digest which is typically smaller than the original reference, SPUMONI 2 builds an *r*-index over the minimizer digest which in turns leads to a smaller index.

Finally, SPUMONI 2 includes a new *sampled document array* structure, allowing for multi-class classification by relating the runs in the *r*-index to a representative document in the input collection.

The user provides a list of genomes along with a class assignment for each at construction time. The sampled document array is small – growing linearly with the size of the index – but allows the user to infer class membership during the computation of matching statistics. Though the sparsity of the structure adds uncertainty to these assignments, this can be compensated for by aggregating assignments along the read.

In short, SPUMONI 2 is a new method and software tool that creates small pangenomic indexes from large collections of genomes. SPUMONI 2 combines the compression of the *r*-index and the adjustable sparsity of minimizers to greatly reduce the index size; for example, indexing 10 human genome sequences along with their reverse complements in 4.2 GB. The classification test used by SPUMONI 2 yields high binary classification accuracy on both short and long reads. Additionally, SPUMONI 2 enables multi-class classification through a new *sampled document array* structure that scales linearly with the amount of distinct sequence. Finally, when comparing SPUMONI 2 to a minimap2-based approach for ONT adaptive sampling in a mock community scenario, SPUMONI 2 is 15 times faster using an index more than 68 times smaller.

## 2 Results

### 2.1 Method Overview

Like SPUMONI, SPUMONI 2 classifies reads according to the lengths of the matching statistics (MSs) computed with respect to an index of reference sequences. In particular, SPUMONI 2 computes an empirical distribution of “null” matching statistics at index construction time, identifying the longest null MS occurring at least a certain number [five] of times.

SPUMONI 2 divides the read into non-overlapping windows of 150 symbols each, starting at the left-end of the read, and classifies the read as “present” in the database if a majority of the windows have a maximum MS longer than the null threshold. As with SPUMONI, SPUMONI 2’s default mode actually computes pseudomatching lengths (PMLs), a fast approximation of matching statistics (MSs) that have similar discriminatory power. Further details are given in Methods (Section 4).

### 2.2 Minimizer digestion

To assess the impact of minimizer digestion, we measured the size of the SPUMONI 2 index when indexing a collection of *Eschericha coli* strains using two minimizer schemes and two alphabets. We chose to index 500 *E. coli* strains from the RefSeq database (24). ^1^ A minimizer scheme is parameterized by a short window size (*k*) and a long window size (*w*). We assessed the (*k, w*)-pairs of (4, 8) and (4, 16). The pair (4, 8) was chosen to yield a digested index with a similar size (*r*) compared to the undigested index. The pair (4, 16) was chosen to allow us to measure the impact of minimizer sparsity on index size. We assessed two choices of alphabet: a minimizer alphabet, where each possible 4-mer is a distinct alphabet symbol, and a typical DNA alphabet with 4 nucleotide symbols. The digested reference was obtained by scanning the reference sequence, computing the minimizers, collapsing equal-minimizer runs, then concatenating the minimizers, either leaving them in the original DNA alphabet or converting them to the minimizer alphabet in the process (Figure 1). We measured both the length of the digested reference (*n*) and the number of runs in its Burrows-Wheeler Transform (*r*), which is the main determinant of index size.

We observed that the compressed indexes of the digested sequences continued to have large compression ratio (*n/r*), indicating that compressed indexing is still effective on minimizer-digested strings (Table 1). In the case of the (4, 8) minimizer scheme, the value of *n* after digestion was greater than than the *n* before digestion (6.78 versus 5.13 billion), but digestion had little effect on *r* (64.4 after digestion versus 65.2 million before). Consistent with expectations, index size was smaller for sparser minimizer schemes.

**Table 1:** SPUMONI index measurements when built over 500 *E. coli* strains (total size 2.5 GB) using different minimizer schemes and alphabets. Both minimizer schemes used a small window size (*k*) of 4.

| | w | No digestion | $w = 8$ | | $w = 16$ | |
| --- | --- | --- | --- | --- | --- | --- |
| alphabet |  |  | Minimizer | DNA | Minimizer | DNA |
| n |  | 5.13 billion | 1.61 billion | 6.78 billion | 663 million | 2.85 billion |
| r |  | 65.2 million | 31.8 million | 64.4 million | 17.5 million | 35.1 million |
| n/r |  | 78.753 | 50.603 | 105.35 | 37.888 | 81.251 |
| index size |  | 232 MB | 142 MB | 227 MB | 73 MB | 124 MB |

As seen in Figure 2a, SPUMONI 2’s index decreased in size as *w* increased. Also, query speed-up was nearly always greater than 1 (i.e. faster than the original index) and also increased with *w*, though with some variability. The minimizer-alphabet indexes (green) always outperformed DNA-alphabet indexes (red) with respect to both index size and speed-up.

**Figure 2:**
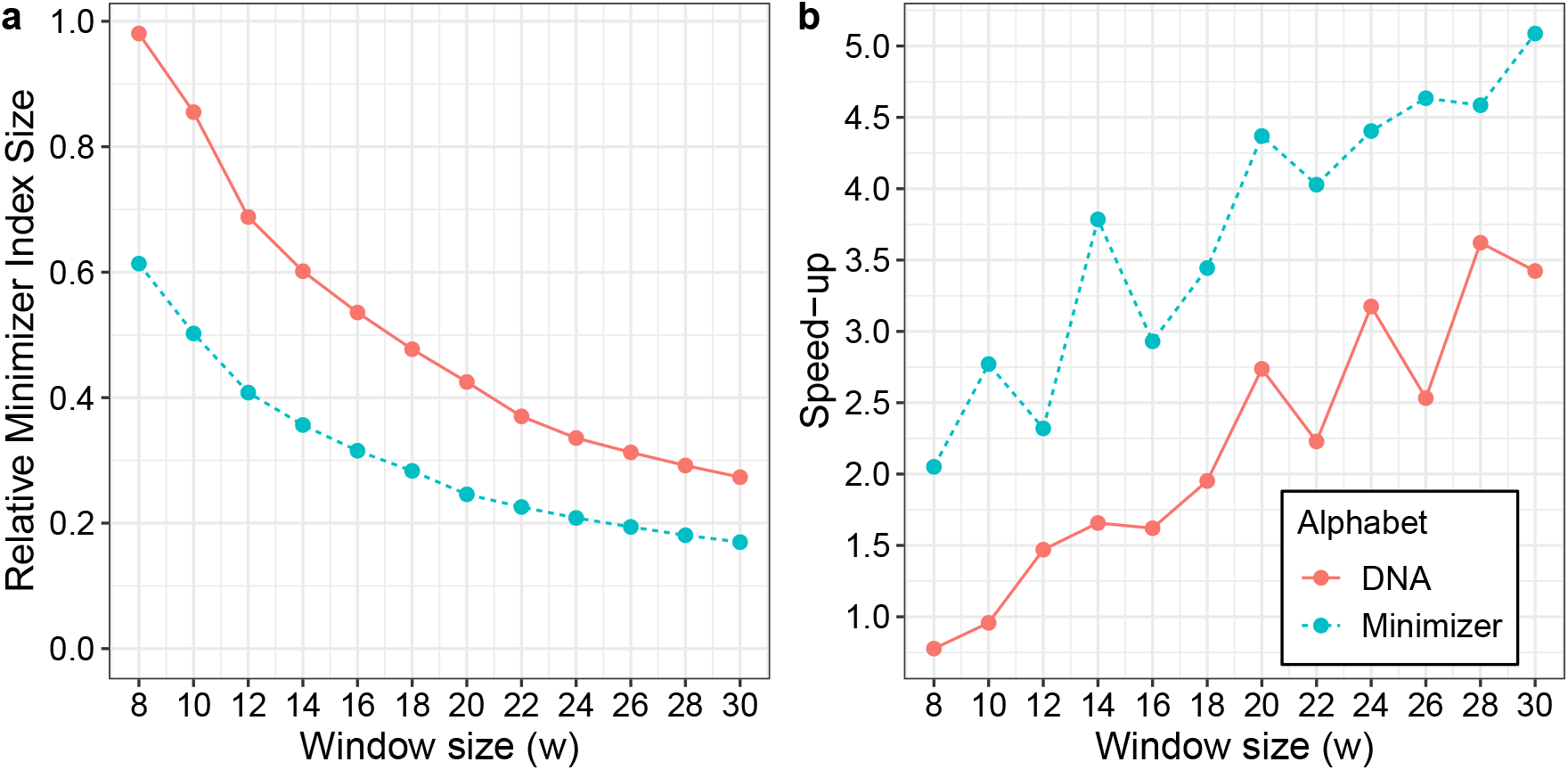
(a) Shows the relative size of the minimizer-based SPUMONI indexes across a range of large window sizes compared to the index size when indexing the full FASTA file. The dataset indexed is a set of 500 *Escherichia coli* genomes, and the small window size was kept at 4. (b) Shows the speed-up achieved when using the minimizer-based indexes to query 1 million short E. coli reads (11) against our index compared to querying against an index over the full FASTA file.

### 2.3 Efficient short and long-read binary classification

To assess how minimizer digestion impacts classification accuracy, we ran SPUMONI 2 with various large window sizes (*w*), using either DNA or minimizer alphabets. We found that the minimizer alphabet allowed SPUMONI 2 to attain high accuracy – as high as the highest accuracy achieved by the DNA alphabet – while yielding a smaller index (Figure 3). While accuracy decreased as the minimizer scheme became sparser (i.e. moving from right to left on the plot), the decrease was less drastic for long reads compared to short reads. For both short and long reads, we setting *w* to a value between 10 and 12 decreased index size with only a minor impact on accuracy. Given these results, we set SPUMONI 2’s default minimizer scheme to use *w* = 11, which will be used in future results.

**Figure 3:**
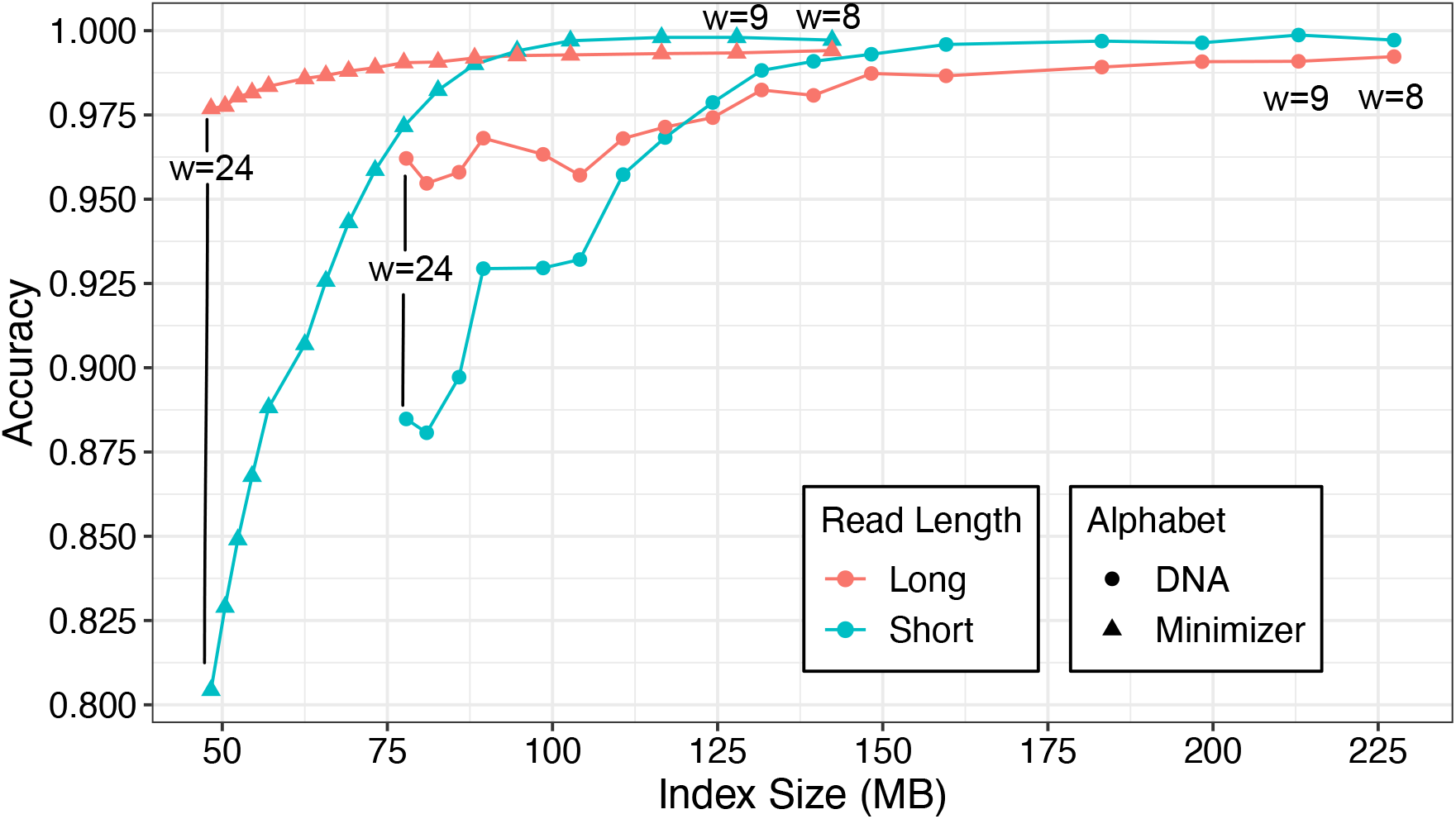
Shows SPUMONI’s binary classification accuracy for indexes of different sizes using different minimizer types. The index was built over 500 *Escherichia coli* genomes and used minimizer schemes where the large window size ranged from 8 to 24. The read set consisted of simulated ONT (mean length = 9000 bp, 95% accuracy) (25) and Illumina reads (150 bp, 99% accuracy) (11) from *E. coli* and Human. The goal was to classify whether the read was from *E. coli* or Human.

### 2.4 Adaptive sampling classification using SPUMONI 2

As a more challenging test case for minimizer digestion, we assessed SPUMONI 2’s classification accuracy in an adaptive sampling setting. More specifically, we replicated the mock-community simulation experiment from the original SPUMONI study (1). We compared SPUMONI to a minimap2-based approach similar to Readfish (26). Classification tools were run on four “batches” consisting of non-overlapping windows of 180 bp from the reads, starting from the read’s left end. Once the tool can make a classification decision, typically after examining the first or second batch, we record the decision and assess accuracy. Experimental details are given in methods (Section 4).

SPUMONI 2 was able to index both pangenomic databases in about half the space as SPUMONI 1, and achieved a 2-fold speed-up with respect to classification time (Table 2). Because we also measured performance of SPUMONI 2 with minimizer digestion disabled (labeled SPUMONI 2**), we can conclude that these improvements were due to minimizer digestion. Compared to minimap2 on the Mock Community dataset, SPUMONI 2 was about 15 times faster with an index more than 68 times smaller (0.74 GB versus 50.9 GB). On the Human Microbiome dataset, the SPUMONI 2’s index was about 15 times smaller than minimap2’s, and SPUMONI 2 was about 4 times faster at classification. The SPUMONI 2 pangenome index comprising of 10 human genome sequences and their reverse complements fit in about 4.2 GB.

**Table 2:** Adaptive sampling simulation using SPUMONI 1, SPUMONI 2 and minimap2. SPUMONI 1 indexes the full input database, while SPUMONI 2 indexes the minimizer-digested sequences of the database using the minimizer alphabet. The “SPUMONI 2 **” gives measurements for SPUMONI 2 with minimizer digestion disabled. Batches of 180 bp (0.4s) of data are delivered in each batch, and the goal is to decide whether to eject the read or not. Four batches were considered in the analysis which corresponds to 720 bp. The mock community dataset of ONT reads (SRX7711546) consists of reads from 7 microbial species and 1 yeast species. The goal is to retain the yeast reads and eject the microbial reads. For the human microbiome study, bacterial reads from the microbiome were obtained the following SRA accession (SRX6602475) and human reads were simulated (25) from the CHM13 reference.

| scenario | Mock Community |  |  |  | Human Microbiome |  |  |  |
| --- | --- | --- | --- | --- | --- | --- | --- | --- |
| goal | retain yeast, eject microbial |  |  |  | retain microbial, eject human |  |  |  |
| index database | 7 microbial species (n=5867 genomes) |  |  |  | human (n=10 genomes) |  |  |  |
| tool | SPUMONI 1 | SPUMONI 2** | SPUMONI 2 | minimap2 | SPUMONI 1 | SPUMONI 2** | SPUMONI 2 | minimap2 |
| sensitivity | 97.38 | 97.62 | 95.32 | 97.90 | 99.24 | 95.07 | 97.08 | 99.56 |
| specificity | 90.77 | 97.38 | 97.06 | 97.85 | 94.30 | 99.97 | 99.12 | 99.97 |
| index size | 1.54 GB | 1.54 GB | 0.74 GB | 50.9 GB | 10.2 GB | 10.2 GB | 4.21 GB | 65.5 GB |
| peak memory | 1.62 GB | 1.62 GB | 0.80 GB | 8.40 GB | 11.0 GB | 11.0 GB | 4.56 GB | 9.99 GB |
| time (s) | 367.39 | 362.89 | 193.53 | 2957.56 | 1628.7 | 1747.12 | 732.63 | 3070.0 |

\*\* running SPUMONI 2 without minimizer digestion (i.e. similar to SPUMONI 1 but using new classification approach)

### 2.5 Contamination detection in assemblies

We expected that SPUMONI 2 would be able to quickly scan for the presence of contaminants in human genome assemblies. NCBI, which curates the GenBank and RefSeq databases, uses a series of filtering protocols to identify contamination, e.g. by using using BLAST (4) to align contigs against common contaminants like *Eschericha coli* and yeast.

We built the SPUMONI 2 index over a pangenome of *Eschericha coli* and yeast genomes (Table S1). The SPUMONI 2 index was 104 MB, 24 times smaller than the total size of the input FASTA files (2.5 GB). We used the contigs from human assemblies as input to SPUMONI 2 to identify contigs with long pseudomatching lengths (PMLs) with respect to the contaminant pangenome. SPUMONI 2 found four contigs in one human assembly (32) that had an average PML of 2 or greater (Figure 4a). When comparing these four to all the other contigs, we observed a stark difference in distribution of PMLs. Figure 4b uses minimap2 alignments to confirm that these contigs have large substrings matching with high identity to the contaminant database. These were reported to the authors of the assembly (32), who verified these findings and removed the contigs from the v2.0 assembly as of December 17, 2021.

**Figure 4:**
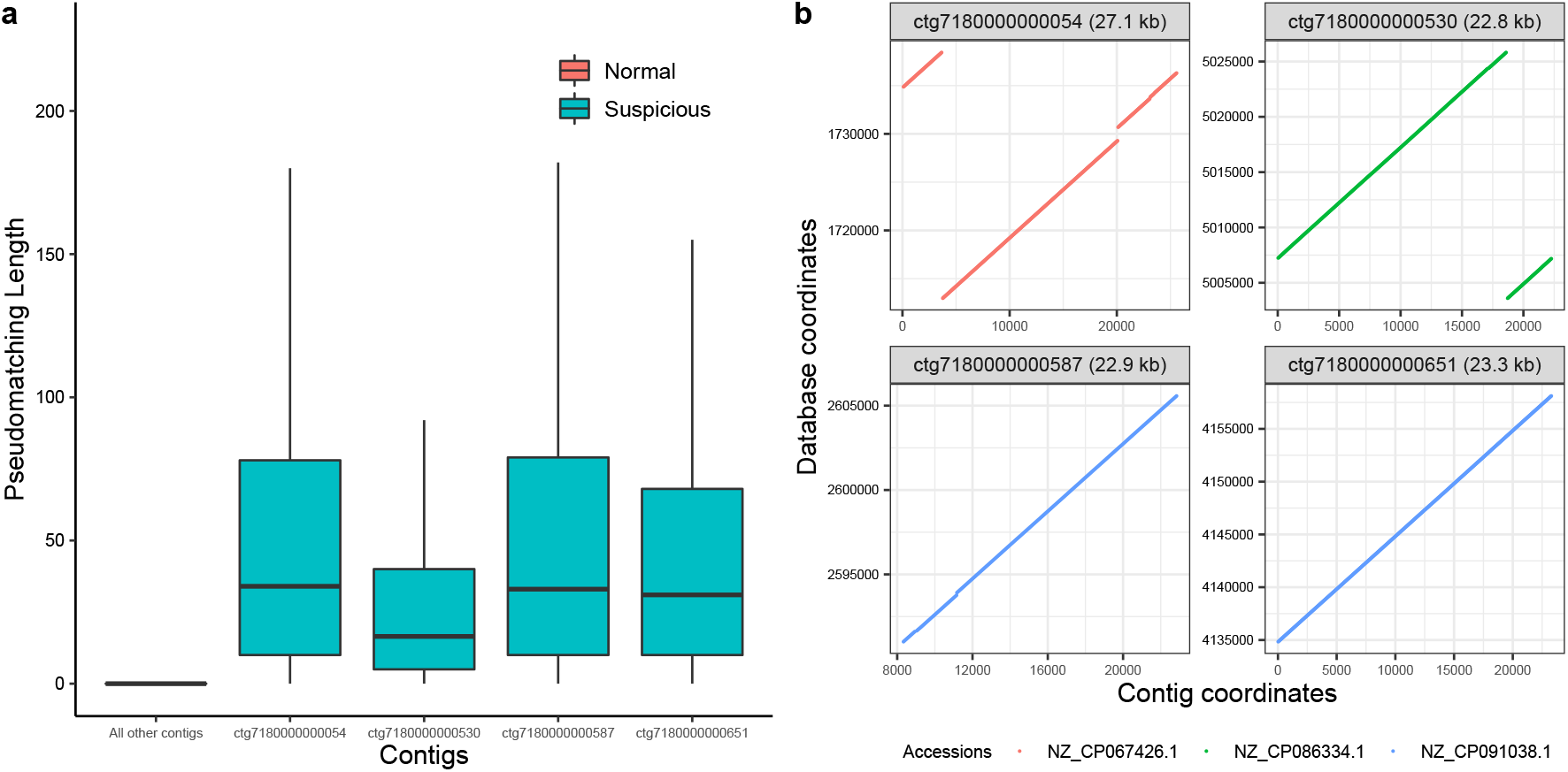
(a) Box plot of pseudomatching lengths (PMLs) for each contig in a human assembly (32) with respect to a SPUMONI 2 index of contaminants. Contigs with an average PML of 2 or greater are labeled “Suspicious.” Contigs with average PML less than 2 are grouped labeled “Normal.”(b) dot plots showing high-scoring local alignments found with minimap2 (18) for the four suspicious contigs versus sequences in the contaminant pangenome. The suspicious contigs were reported and moved from the assembly in December, 2021

We compared SPUMONI 2’s speed to that of BLAST+, which is used by NCBI for assembly filtering. SPUMONI 2 was 4.3 times faster while identifying the same suspicious contigs (Table S3).

### 2.6 Small multi-class classification using the sampled document array

We assessed SPUMONI 2’s sampled document array in a multi-class classification setting. As SPUMONI 2 computes matching statistics at each position of the read, it queries the sampled document array to obtain a class label for one document (sequence) containing the current match. The process involves uncertainty, since the particular class reported by the sampled document array may be just one among many that contain the match (see Section 4.7 for details). We hypothesized that, by aggregating the classes reported over the course of all the steps of the algorithm, we can infer the correct class of origin.

We used SPUMONI 2 to construct a pangenomic index over all of the genomes in RefSeq for eight different microbial species (details in Table S2). We observed that the sampled document array increased the size of the index from 793 MB to 920 MB, a 16% increase.

Figure 5 shows that when querying simulated Illumina reads (11) from eight different microbial species against a pangenomic database, we find that between 80.7% and 88.0% of the classes reported by the sampled document array at the read level match the true class of origin. For long ONT reads, we see a similar pattern where the true class is in the majority, ranging from 53.8% to 66.3% of the labels across the eight classes (Figure S1).

**Figure 5:**
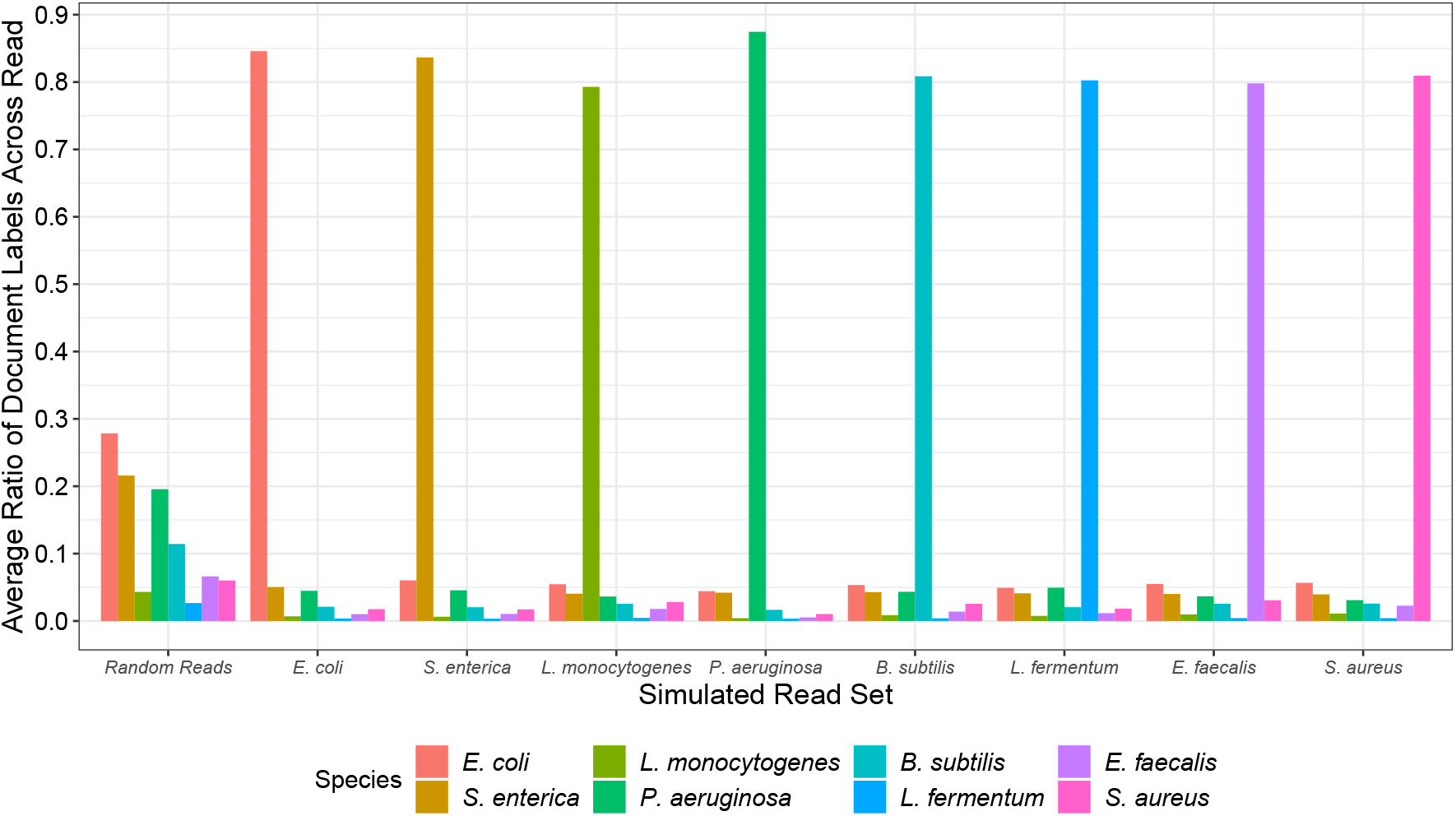
Average ratio of class labels found at the read level when matching reads that were simulated from eight microbial species. The SPUMONI 2 index was a pan-genome database of microbial species.

When queried with randomly-generated reads consisting of independent and uniform draws from the DNA alphabet, no class label exceeded 28.2% frequency for short reads (Figure 5). We observed a tendency for some classes to be reported more often than others (e.g. *E. coli*), likely because of differences in the total number of indexed bases of each class (Table S2).

## 3 Discussion

SPUMONI 2 is a read classification tool optimized for classifying both short and long reads against a pangenomic database. By combining minimizer digestion with the *r*-index, SPUMONI 2 compresses the pangenome more efficiently than SPUMONI, allowing it to handle larger databases. Compared to minimap2, SPUMONI 2’s index is far smaller (1/68th of minimap2’s in the case of our mock community experiment) and it performs classification up to 15 times faster. While SPUMONI 2’s computational efficiency comes at the expense of some classification accuracy, this can be a favorable trade in situations where the pangenome is very large (e.g. minimap2’s index of 10 human genome assemblies is over 50 GB) or where reaction time is important, e.g., in an adaptive sampling setting where the algorithm has to keep up with DNA strings moving through many hundreds of pores simultaneously.

SPUMONI 2 includes a novel sampled document array structure whose space usage achieves the same *O*(*r*) bound as the overall *r*-index. This allows our matching-statistics-based classification method to work in multi-class classification settings, expanding its applicability to metagenomics classification.

While the sampled document array was effective in the long-read classification scenario studied here, it is limited by the fact that it records only one “document” (class) per *r*-index run boundary. A more general approach would additionally allow us to query all classes containing the current match, not just the class corresponding to the run boundary. Such approaches exist for uncompressed indexes, but more work is needed to adapt these to work in compressed space and time. This will be particularly important for short-read classification, where we cannot aggregate as many document array queries.

Finally, we demonstrated a novel application of SPUMONI 2 to detecting contaminants in genome assemblies. This represents another application where, as reference databases continue to grow, SPUMONI 2 will be well positioned to leverage the additional assemblies with minimal impact to index size or classification speed. On the other hand, larger genomic databases have been shown to affect *k*-mer based approaches due to decreasing levels of *k*-mer specificity (22).

## 4 Methods

### 4.1 Compressed indexes

Given a text *T* [1..*n*] of length *n*, the Burrows-Wheeler transform (BWT) (3) is a reversible permutation of *T*’s characters such that BWT[*i*] is the character preceding the *i*^th^ lexicographical suffix of *T*. The BWT of a repetitive text will have long runs of the same character. The symbol *r* denotes the number of maximal same-character runs in the BWT. The *r*-index (10) is a compressed index consisting of the run-length encoded BWT, plus auxiliary structures enabling rapid queries for counting the number of times a substring occurs in *T*, as well as for locating all offsets in *T* where a substring occurs. Only *O*(*r*) space is needed overall. More details can be found in the study of Gagie et al (9).

### 4.2 Matching statistics and MONI

The matching statistics array MS[1..*m*] is defined for a pattern *P* [1..*m*] with respect to a text *T* [1..*n*]. The element MS[*i*] equals the length of the longest prefix of *P*’s *i*^th^ suffix that occurs in *T*. Using the *r*-index, Bannai et al. (2) proposed a two-pass algorithm for computing MS using an additional “thresholds” structure. The algorithm’s first pass iterates over *P* from right to left, attempting to use the *LF* mapping at each step to extend the match to the left by one character. If the algorithm reaches a row *j* and finds that match cannot be extended because the next character of *P* mismatches BWT[*j*], the algorithm skips (“jumps”) either up or down to the next BWT run that does start with the next character of *P*. The direction of the jump is determined by the threshold, which indicates whether jumping up or down yields a longer match. Given the steps taken through the BWT in the first pass of the algorithm, the second pass uses a random-access data-structure over *T* to compute the exact matching statistic lengths for MS. More details can be found in the study of Rossi et al (29).

### 4.3 Pseudomatching lengths and SPUMONI

Pseudomatching lengths (PMLs) were proposed in the SPUMONI study as being a quantity similar to matching statistics but more efficient to compute. SPUMONI’s computation of PMLs does not require the second pass of the algorithm described above. Instead, each step of the first pass that successfully extends the match using the LF mapping causes a length variable to be incremented by one. When the algorithm reaches a step where it cannot proceed using the LF mapping and has to jump instead, the length variable is reset to 0. The values taken by this variable at each step constitutes the vector of PMLs (PML). The PML values are upper-bounded by the MS values, but we have shown that they are similarly useful for classification of sequences (1).

The SPUMONI index consists chiefly of the *r*-index (omitting the sampled suffix array) and the thresholds. These structures allow SPUMONI to compute PML in *O*(*r*) space. Because the *r*-index component does not need to include the sampled suffix array, SPUMONI’s disk and memory footprint can be substantially smaller than that of the MONI algorithm described in the previous subsection.

### 4.4 Minimizer digestion

SPUMONI 2 computes minimizers by applying a hash function to all *k*-mers in a larger window of size *w, w > k*. The *k*-mer with minimal hash value is the minimzer. If the minimizers found in two or more consecutive steps are identical, they are compressed to a single copy of the minimizer. SPUMONI 2 uses the RollingHasher object provided by the Bonsai C++ library.

These minimizer sequences are then indexed by using the *r*-index (9) to build a runlength encoded BWT and thresholds data-structure.

### 4.5 Read classification

The previous SPUMONI method classified reads by accumulating an empirical distribution of positive PMLs (with respect to the reference), and another of null PMLs (with respect to the reverse of the reference) (1). It used a Kolmogorov-Smirnov statistical test (KS-test) to assess whether the distribution of positive PMLs was overall larger (shifted higher) compared to the null PMLs.

SPUMONI 2 uses a simpler approach; it also accumulates empirical distributions of positive and null PMLs, but it classifies reads by first computing a threshold PML value as a function of the null PML distribution. The null PML distribution is constructed by extracting small reads (substrings) from the reference, reversing the sequence, then computing PMLs for those sequences with respect to the reference. Since the sequences are reversed (not reverse complemented), they serve as random sequences that should not be classified as matching the reference, but they nonetheless share the reference’s base distribution. Pooling across simulated reads, we compile an aggregate distribution of null PMLs and compute the threshold as being equal to the largest PML that occurs at least 5 times. Because the process of computing the threshold uses the same parameters as the read-classification process (i.e. same minimizer scheme and alphabet), the choice of threshold is customized to the problem at hand.

Given the threshold PML value, SPUMONI 2 classifies a read by computing the read’s positive PMLs with respect to the reference index in many distinct non-overlapping windows of the read. By default, the read is divided into non-overlapping windows of 150 symbols. We found this window size performs well in terms of accuracy across different alphabets (minimizers or DNA). The windows at the end of the read that are less than 150 symbols are grouped together with the previous window. If a majority of the windows have a maximum PML greater than the threshold, the read is classified as matching the reference, otherwise it is classified as not matching.

### 4.6 Adaptive sampling simulation

Nanopore sequencers report a time series representing the amount of current flowing through a pore. The data is split into batches which are delivered to the control software for analysis. Following our experimental setup from the SPUMONI study (1), we simulated 4 batches of 0.4 sec of basecalled nanopore data, with each batch consisting of 180 bp each. Batches of this size were shown to be sufficient for binary classification method (18, 26). We start from base-called data, rather than from raw signal data.

For our adaptive sampling simulation in Section 2.4, we simulated two different scenarios where adaptive sampling could be used to enrich for certain reads. The first scenario is a mock community which consists seven bacterial species (*Staphylococcus aureus, Salmonella enterica, Escherichia coli, Pseudomonas aeruginosa, Listeria monocytogenes, Enterococcus faecalis, Bacillus subtilis*) and 1 yeast species (*Saccharomyces cerevisiae*). The goal in this scenario is to enrich for the yeast reads by rejecting bacterial reads. To accomplish this, we built a pangenome reference over all the strains of the seven bacterial species in RefSeq (24) to identify which reads come from bacteria.

The second scenario is focused on the human microbiome where we used a 50:50 mixture of simulated human reads from the CHM13 reference (23), and real nanopore reads from the microbial species in the human gut (21). The goal in this experiment is to enrich for the microbial reads while removing as many human reads as possible. For this scenario, we built a pangenome reference over 10 human assemblies (5, 6, 12, 17, 23, 27, 31, 33, 35) in order to identify human reads that we want to remove.

We use both SPUMONI and minimap2 to classify batches against the pangenome index. A read classified as “present” should be immediately ejected by the pore, so subsequent batches of data are not processed. SPUMONI 2 classifies the current batch of 180 bp using the test described in section (4.5). In the case of minimap2, aligning against a pangenomic database usually yields numerous alignments: a primary and many secondary alignments. We examine these to ensure all are to the same species and, if so, we classify the read as “present.” For any batch after the first, we provide the entire base-called read so far to minimap2; i.e. we concatenate the bases from all the batches so far. In this way, minimap2 is redundantly processing the same bases when examining batches beyond the first. This is consistent with minimap2’s design; unlike the *r*-index-based algorithms that have the ability to “pause” and “resume” the matching process, minimap2 must be run on a complete read sequence.

The time required for each method is reported in Table 2; this is the total time used by each method to classify the reads across all 4 batches of data.

### 4.7 Sampled document array

SPUMONI 2’s sampled document array consists of 2*r* integers encoding the class of the suffixes at the beginning and end of every BWT run. The class labels are computed at index construction time. Note that the class labels can be stored in ⌈log_2_ *c*⌉ bits, where *c* is the number of class labels, which willl often be much less than the ⌈log_2_ *n*⌉ bits required for suffix array entries. An example is shown in Figure S4.

At query time, the sampled document array is queried whenever the algorithm for computing PMLs passes through a suffix at a run boundary. As described in detail in the MONI paper (29), the algorithm proceeds base-by-base starting from the right-hand extreme of the read. Each step falls into one of two cases. When the match can be extended to the left using the LF mapping, this is called “case 1.” When the match cannot be extended to the left using the LF mapping, the algorithm uses threshold information to choose a new BWT run to “jump” to. This is “case 2.” Importantly, case 2 always results in the algorithm moving to a run boundary. That is, when the algorithm jumps, it jumps either to the beginning of a run (if it jumps down) or the end of a run (if it jumps up).

When using the sampled document array, the SPUMONI 2 algorithm compiles not only its usual array of PMLs, but also an array of document (class) labels, CA. At position *j* in the read where the algorithm uses case 2, the label CA[*j*] equals the sampled document array element corresponding to the run boundary jumped to. For a position *j* in the read where the algorithm uses case 1, the label CA[*j*] equals the class observed in the most recent instance of case 2. When the read truly originates from one of the classes in the reference pangenome, we expect the majority of the labels in CA to match the true class. This algorithm is summarized in the form of pseudocode in Supplementary Figure S2.

## 5 Acknowledgments

This work was carried out at the Advanced Research Computing at Hopkins (ARCH) core facility (rockfish.jhu.edu), which is supported by the National Science Foundation (NSF) grant number OAC 1920103.

## 7 Availability of data and materials

- The SPUMONI 2 software is available from GitHub at https://github.com/oma219/spumoni
- The code used to run the experiments is available at https://github.com/oma219/spumoni-experiments
- FASTA files for the E. coli genomes assessed in sections 2.2 and 2.3 are available at https://genome-idx.s3.amazonaws.com/spu2/ecoli_500_dataset.tar.gz
- SPUMONI 2 index files for the mock community pangenomes assessed in section 2.4 are available at https://genome-idx.s3.amazonaws.com/spu2/mock_community_ont_index.tar.gz
- SPUMONI 2 index files for the human assembly pangenome assessed in section 2.4 are available at https://genome-idx.s3.amazonaws.com/spu2/mock_community_ont_index.tar.gz
- SPUMONI 2 index files for the assembly contaminant pangenome used in section 2.5 are available at https://genome-idx.s3.amazonaws.com/spu2/assembly_contamination_index.tar.gz
- SPUMONI 2 index files, including the sampled document array, for the experiments described in section 2.6 are available at https://genome-idx.s3.amazonaws.com/spu2/sampled_doc_array_index.tar.gz

## 8 Authors’ contributions

OA and BL conceived the project and designed SPUMONI 2. OA implemented the SPUMONI 2 software with assistance from BL and MR. All authors contributed to the design of the sampled document array structure. All authors wrote, edited and approved the manuscript.

## 10 Additional Files

## Supplementary Notes

### Read Simulation

In the paper, read simulators such as Mason and PBSIM2 were used to simulate short and long reads respectively. The commands used to generate those reads are shown below:

~~~
   mason_simulator -ir $genome -n $num_of_reads -v \
                   -o $output_read --illumina-read-length 150 \
                   --illumina-prob-mismatch 0.01
~~~

This command above was generally used throughout the paper for simulating short reads with 1% mismatch error. Next, the command below shows how long reads were simulated with 95% read accuracy using R9.4 chemistry.

~~~
  pbsim --depth 50.0 --prefix $output_prefix \
        --hmm_model R9.4.model --accuracy-mean 0.95 \
        $positive_genome
~~~

## Supplementary Figures

**Figure 1:**
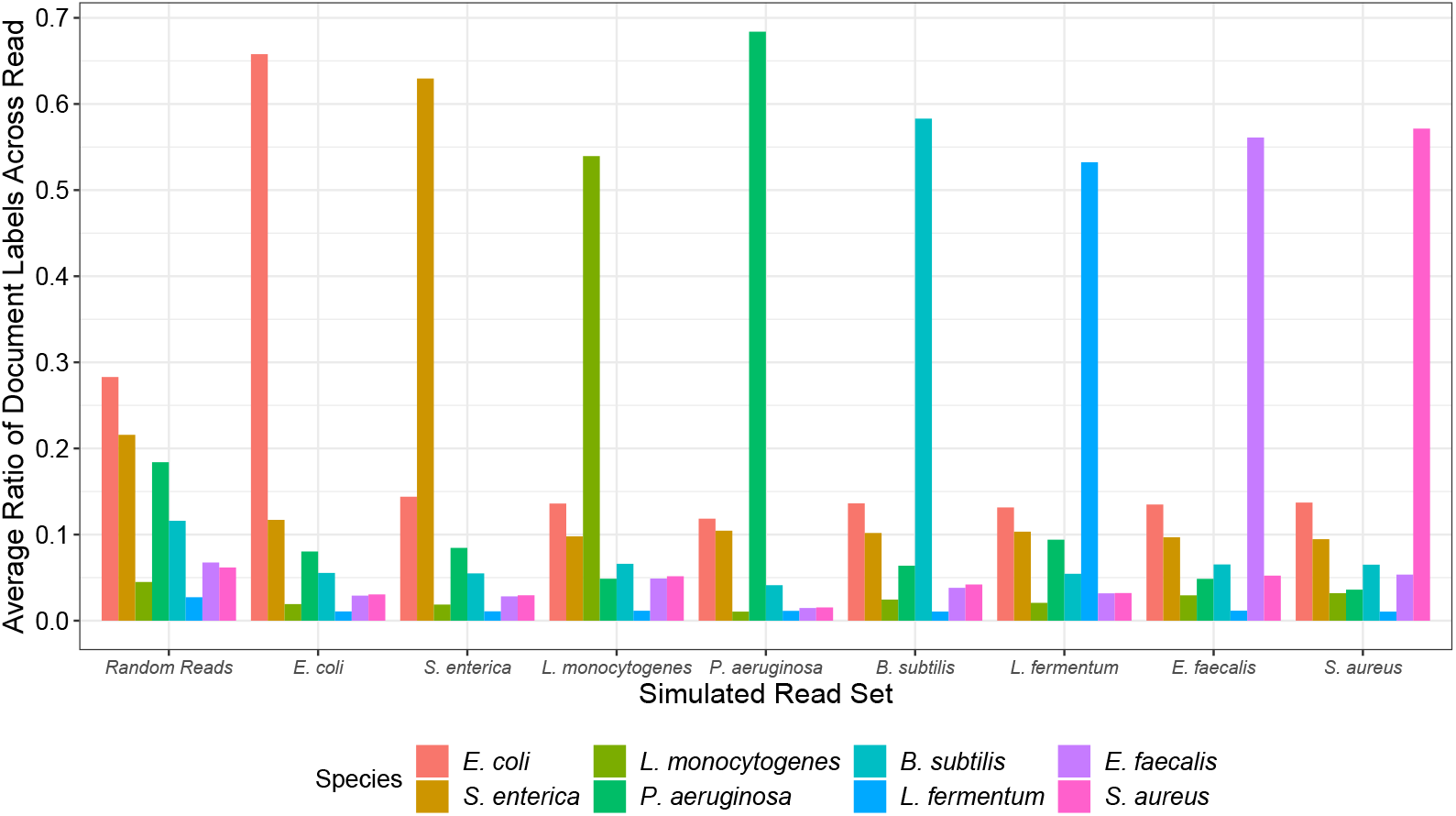
Shows the average ratio of document labels found at the read level when querying simulated long reads (ONT) from eight different microbial species against a pangenome database of microbial species.

**Figure 2:**
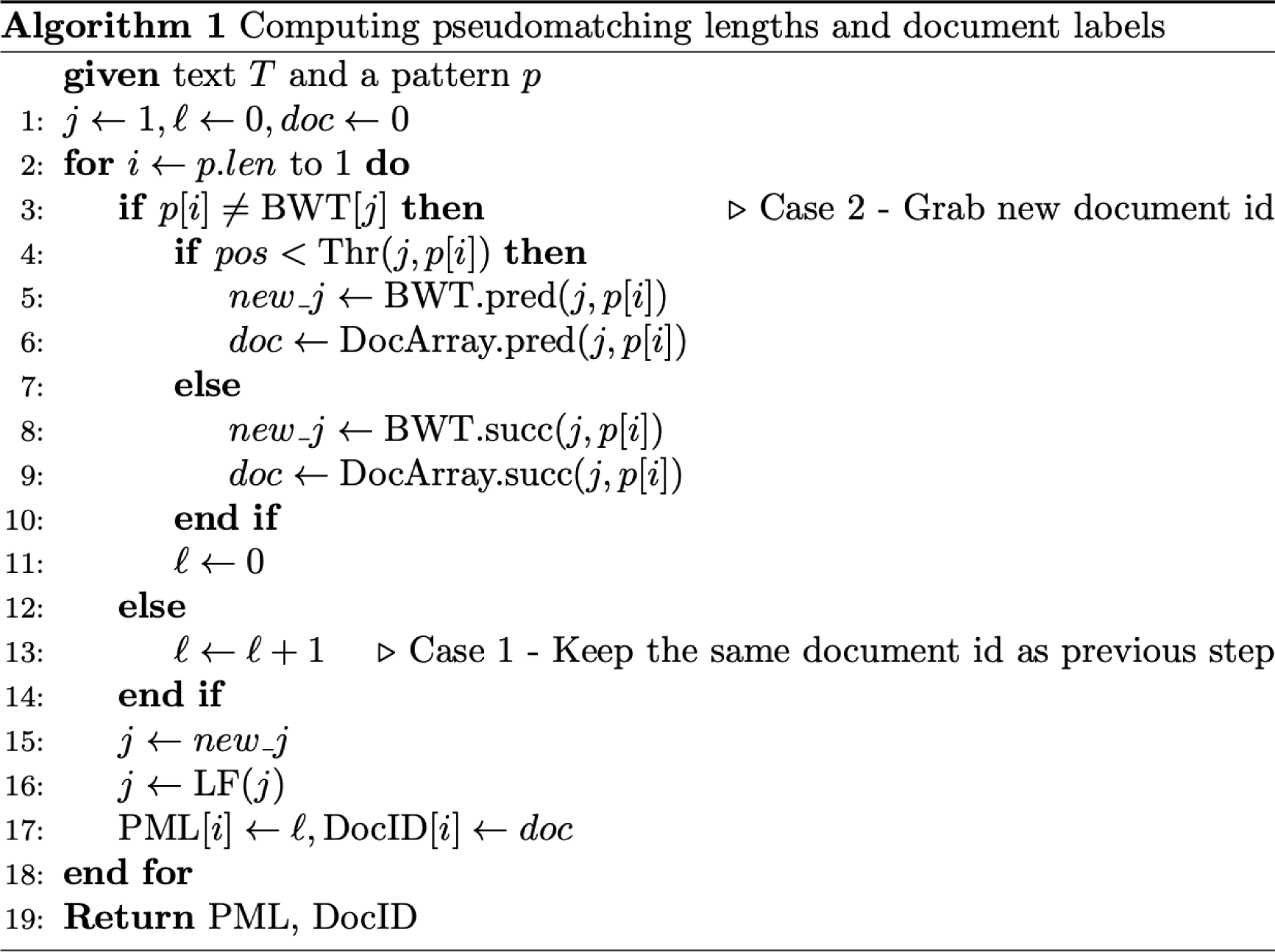
Shows the pseudocode for computing pseudomatching lengths as well as document labels for a pattern *P* with respect to a text *T*.

## Supplementary Tables

**Table 1:** Shows the the different species and the number of genomes used in the “contaminant” database for the assembly contamination experiment. All of the strains for the species listed above were obtained from the NCBI RefSeq database.

| “Contaminant” Database Contents |  |
| --- | --- |
| Species: | Number of Genomes: |
| *Escherichia Coli* | 500 |
| *Saccharomyces cerevisiae* | 1 |

**Table 2:** Shows the eight microbial species used in the document array experiment, and the number of genomes from RefSeq that were included in the database. All of the strains for the species listed above were obtained from the NCBI RefSeq database.

| Pan-genomic database for small multi-class classification experiment |  |  |
| --- | --- | --- |
| Species: | Number of Genomes: | Number of Bases: |
| *Escherichia Coli* | 2495 | 12.9 billion |
| *Salmonella enterica* | 1309 | 6.40 billion |
| *Listeria monocytogenes* | 281 | 833 million |
| *Pseudomonas aeruginosa* | 470 | 3.15 billion |
| *Bacillus subtilis* | 217 | 899 million |
| *Lactobacillus fermentum* | 40 | 85.2 million |
| *Enterococcus faecalis* | 459 | 1.38 billion |
| *Staphylococcus aureus* | 861 | 2.44 billion |

**Table 3:**
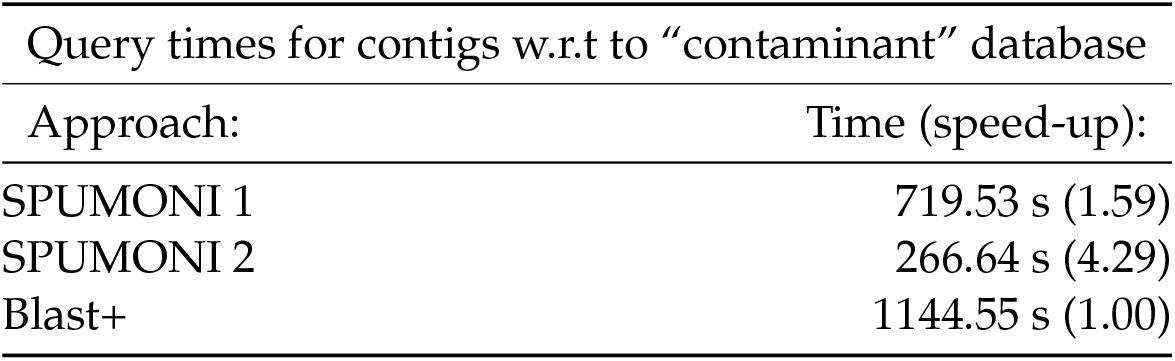
Shows the time taken to query the assembly with respect to “contaminant” database for each method in seconds. Each tool was run with 16 threads. All the tools agreed in terms of which contigs had suspicious similarity to the “contaminant” database.

**Table 4:**
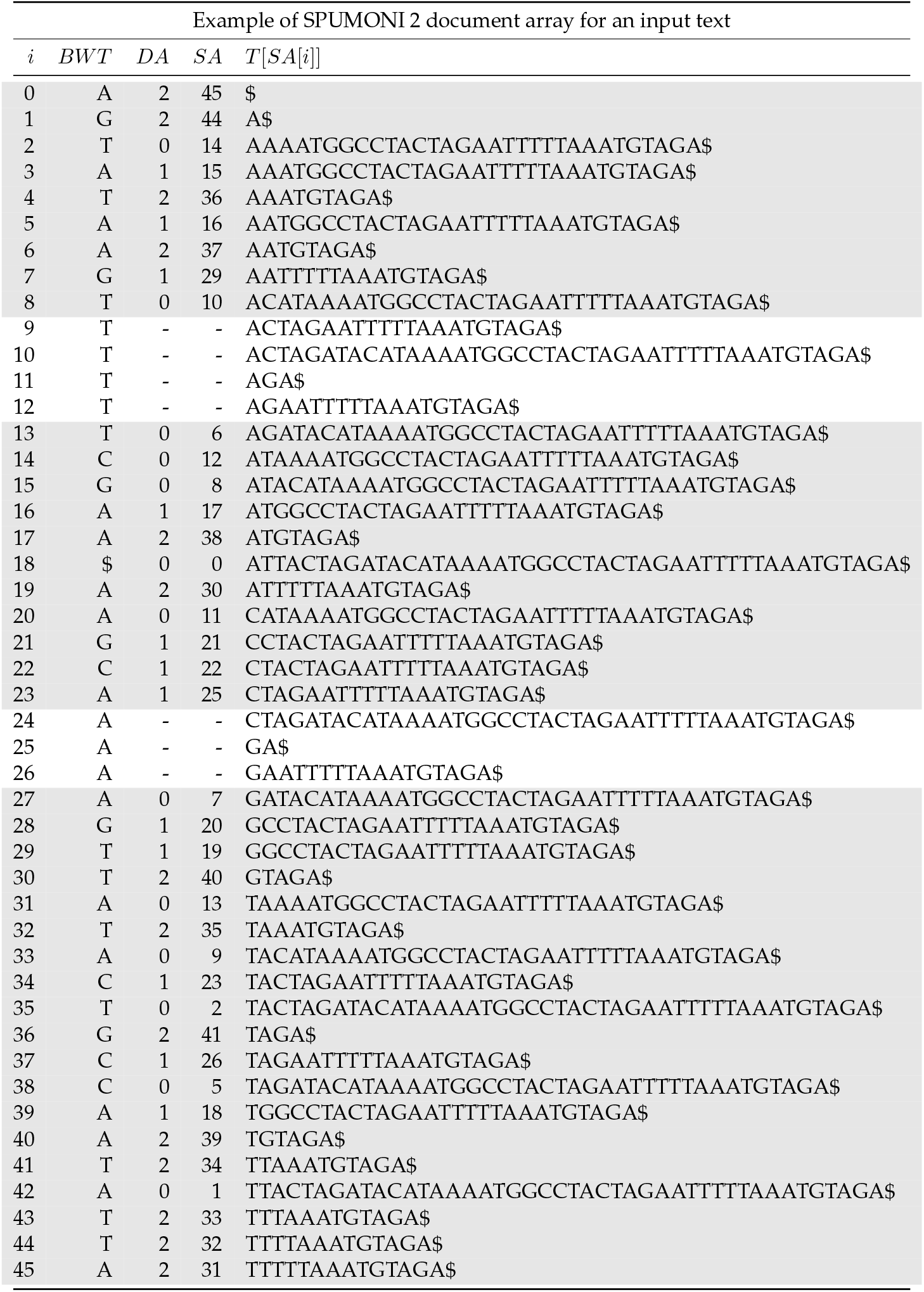
Shows the SPUMONI 2 document array for the collection of documents *T* = {ATTACTAGATACATA, AAATGGCCTACTAGA, ATTTTTAAATGTAGA}. The docu-ment array stores the class number that the suffix at the run boundaries occurs in. The gray rows in the table are labeling the run boundaries of the BWT, which are the ones containing document-array entries. Note that the document array will be much sparser for a larger, more repetitive text.

## Footnotes

1 We give the accessions numbers in the supplemental data.

## References

[1] Ahmed, O., Rossi, M., Kovaka, S., Schatz, M. C., Gagie, T., Boucher, C., and Langmead, B. (2021). Pan-genomic matching statistics for targeted nanopore sequencing. Iscience, 24(6), 102696.

[2] Bannai, H., Gagie, T., and Tomohiro, I. (2020). Refining the r-index. Theoretical Computer Science, 812, 96–108.

[3] Burrows, M. and Wheeler, D. (1994). A block-sorting lossless data compression algorithm. Digital SRC Research Report.

[4] Camacho, C., Coulouris, G., Avagyan, V., Ma, N., Papadopoulos, J., Bealer, K., and Madden, T. L. (2009). Blast+: architecture and applications. BMC bioinformatics, 10(1), 1–9.

[5] Chaisson, M. J., Huddleston, J., Dennis, M. Y., Sudmant, P. H., Malig, M., Hormozdiari, F., Antonacci, F., Surti, U., Sandstrom, R., Boitano, M., et al. (2015). Resolving the complexity of the human genome using single-molecule sequencing. Nature, 517(7536), 608–611.

[6] Church, D. M., Schneider, V. A., Steinberg, K. M., Schatz, M. C., Quinlan, A. R., Chin, C. S., Kitts, P. A., Aken, B., Marth, G. T., Hoffman, M. M., Herrero, J., Mendoza, M. L., Durbin, R., and Flicek, P. (2015). Extending reference assembly models. Genome Biol, 16, 13.

[7] Coombe, L., Nikolić, V., Chu, J., Birol, I., and Warren, R. L. (2020). ntjoin: Fast and lightweight assembly-guided scaffolding using minimizer graphs. Bioinformatics, 36(12), 3885–3887.

[8] Ekim, B., Berger, B., and Chikhi, R. (2021). Minimizer-space de bruijn graphs: Wholegenome assembly of long reads in minutes on a personal computer. Cell systems, 12(10), 958–968.

[9] Gagie, T., Navarro, G., and Prezza, N. (2018). Optimal-time text indexing in bwt-runs bounded space. In Proceedings of the Twenty-Ninth Annual ACM-SIAM Symposium on Discrete Algorithms, pages 1459–1477. SIAM.

[10] Gagie, T., Navarro, G., and Prezza, N. (2020). Fully functional suffix trees and optimal text searching in bwt-runs bounded space. Journal of the ACM (JACM), 67(1), 1–54.

[11] Holtgrewe, M. (2010). Mason: a read simulator for second generation sequencing data.

[12] Huddleston, J., Chaisson, M. J., Steinberg, K. M., Warren, W., Hoekzema, K., Gordon, D., Graves-Lindsay, T. A., Munson, K. M., Kronenberg, Z. N., Vives, L., et al. (2017). Discovery and genotyping of structural variation from long-read haploid genome sequence data. Genome research, 27(5), 677–685.

[13] Kim, D., Song, L., Breitwieser, F. P., and Salzberg, S. L. (2016). Centrifuge: rapid and sensitive classification of metagenomic sequences. Genome research, 26(12), 1721–1729.

[14] Kovaka, S., Fan, Y., Ni, B., Timp, W., and Schatz, M. C. (2021). Targeted nanopore sequencing by real-time mapping of raw electrical signal with uncalled. Nature biotechnology, 39(4), 431–441.

[15] Kuhnle, A., Mun, T., Boucher, C., Gagie, T., Langmead, B., and Manzini, G. (2020). Efficient construction of a complete index for pan-genomics read alignment. Journal of Computational Biology, 27(4), 500–513.

[16] Langmead, B. and Salzberg, S. L. (2012). Fast gapped-read alignment with bowtie 2. Nature methods, 9(4), 357–359.

[17] Levy, S., Sutton, G., Ng, P. C., Feuk, L., Halpern, A. L., Walenz, B. P., Axelrod, N., Huang, J., Kirkness, E. F., Denisov, G., et al. (2007). The diploid genome sequence of an individual human. PLoS biology, 5(10), e254.

[18] Li, H. (2018). Minimap2: pairwise alignment for nucleotide sequences. Bioinformatics, 34(18), 3094–3100.

[19] Li, W., O’Neill, K. R., Haft, D. H., DiCuccio, M., Chetvernin, V., Badretdin, A., Coulouris, G., Chitsaz, F., Derbyshire, M. K., Durkin, A. S., Gonzales, N. R., Gwadz, M., Lanczycki, C. J., Song, J. S., Thanki, N., Wang, J., Yamashita, R. A., Yang, M., Zheng, C., Marchler-Bauer, A., and Thibaud-Nissen, F. (2021). RefSeq: expanding the Prokaryotic Genome Annotation Pipeline reach with protein family model curation. Nucleic Acids Res, 49(D1), D1020–D1028.

[20] Menzel, P., Ng, K. L., and Krogh, A. (2016). Fast and sensitive taxonomic classification for metagenomics with kaiju. Nature communications, 7(1), 1–9.

[21] Moss, E. L., Maghini, D. G., and Bhatt, A. S. (2020). Complete, closed bacterial genomes from microbiomes using nanopore sequencing. Nature Biotechnology, 38(6), 701–707.

[22] Nasko, D. J., Koren, S., Phillippy, A. M., and Treangen, T. J. (2018). Refseq database growth influences the accuracy of k-mer-based lowest common ancestor species identification. Genome biology, 19(1), 1–10.

[23] Nurk, S., Koren, S., Rhie, A., Rautiainen, M., Bzikadze, A. V., Mikheenko, A., Vollger, M. R., Altemose, N., Uralsky, L., Gershman, A., et al. (2022). The complete sequence of a human genome. Science, 376(6588), 44–53.

[24] O’Leary, N. A., Wright, M. W., Brister, J. R., Ciufo, S., Haddad, D., McVeigh, R., Rajput, B., Robbertse, B., Smith-White, B., Ako-Adjei, D., et al. (2016). Reference sequence (refseq) database at ncbi: current status, taxonomic expansion, and functional annotation. Nucleic acids research, 44(D1), D733–D745.

[25] Ono, Y., Asai, K., and Hamada, M. (2021). Pbsim2: a simulator for long-read sequencers with a novel generative model of quality scores. Bioinformatics, 37(5), 589–595.

[26] Payne, A., Holmes, N., Clarke, T., Munro, R., Debebe, B. J., and Loose, M. (2021). Readfish enables targeted nanopore sequencing of gigabase-sized genomes. Nature biotechnology, 39(4), 442–450.

[27] Pendleton, M., Sebra, R., Pang, A. W. C., Ummat, A., Franzen, O., Rausch, T., Stütz, M., Stedman, W., Anantharaman, T., Hastie, A., et al. (2015). Assembly and diploid architecture of an individual human genome via single-molecule technologies. Nature methods, 12(8), 780–786.

[28] Roberts, M., Hayes, W., Hunt, B. R., Mount, S. M., and Yorke, J. A. (2004). Reducing storage requirements for biological sequence comparison. Bioinformatics, 20(18), 3363–3369.

[29] Rossi, M., Oliva, M., Langmead, B., Gagie, T., and Boucher, C. (2022). Moni: A pangenomic index for finding maximal exact matches. Journal of Computational Biology, 29(2), 169–187.

[30] Sayers, E. W., Bolton, E. E., Brister, J. R., Canese, K., Chan, J., Comeau, D. C., Connor, R., Funk, K., Kelly, C., Kim, S., Madej, T., Marchler-Bauer, A., Lanczycki, C., Lathrop, S., Lu, Z., Thibaud-Nissen, F., Murphy, T., Phan, L., Skripchenko, Y., Tse, T., Wang, J., Williams, R., Trawick, B. W., Pruitt, K. D., and Sherry, S. T. (2022). Database resources of the national center for biotechnology information. Nucleic Acids Res, 50(D1), D20–D26.

[31] Seo, J.-S., Rhie, A., Kim, J., Lee, S., Sohn, M.-H., Kim, C.-U., Hastie, A., Cao, H., Yun, J.-Y., Kim, J., et al. (2016). De novo assembly and phasing of a korean human genome. Nature, 538(7624), 243–247.

[32] Shumate, A., Zimin, A. V., Sherman, R. M., Puiu, D., Wagner, J. M., Olson, N. D., Pertea, M., Salit, M. L., Zook, J. M., and Salzberg, S. L. (2020). Assembly and annotation of an ashkenazi human reference genome. Genome biology, 21(1), 1–18.

[33] Steinberg, K. M., Schneider, V. A., Graves-Lindsay, T. A., Fulton, R. S., Agarwala, R., Huddleston, J., Shiryev, S. A., Morgulis, A., Surti, U., Warren, W. C., et al. (2014). Single haplotype assembly of the human genome from a hydatidiform mole. Genome research, 24(12), 2066–2076.

[34] Wood, D. E., Lu, J., and Langmead, B. (2019). Improved metagenomic analysis with kraken 2. Genome biology, 20(1), 1–13.

[35] Zook, J. M., Catoe, D., McDaniel, J., Vang, L., Spies, N., Sidow, A., Weng, Z., Liu, Y., Mason, C. E., Alexander, N., et al. (2016). Extensive sequencing of seven human genomes to characterize benchmark reference materials. Scientific data, 3(1), 1–26.

